# A role for RNA and DNA:RNA hybrids in the modulation of DNA repair by homologous recombination

**DOI:** 10.1101/255976

**Authors:** Giuseppina D’Alessandro, Marek Adamowicz, Donna Whelan, Sean Michael Howard, Corey Winston Jones-Weinert, Valentina Matti, Eli Rothenberg, Petr Cejka, Fabrizio d’Adda di Fagagna

## Abstract

DNA double-strand breaks (DSBs) are toxic DNA lesions which, if not properly repaired, may lead to genomic instability, cell death and senescence. Damage-induced long non-coding RNAs (dilncRNAs) are transcribed from broken DNA ends and contribute to DNA damage response (DDR) signaling. Here we show that dilncRNAs play a role in DSB repair by homologous recombination (HR) by contributing to the recruitment of the HR proteins BRCA1, BRCA2, and RAD51, without affecting DNA-end resection. In S/G2-phase cells, dilncRNAs pair to the resected DNA ends and form DNA:RNA hybrids, which are recognized by BRCA1 and promote its recruitment to DSBs. We also show that RNase H2 is in a complex with the HR proteins BRCA1, PALB2, BRCA2, and RAD51, and that it localizes to DSBs in the S/G2 cell-cycle phase. BRCA2 controls DNA:RNA hybrid levels at DSBs by mediating RNase H2 recruitment and, therefore, hybrids degradation. These results demonstrate that regulated DNA:RNA hybrid levels at DSBs contribute to HR-mediated repair.

## Introduction

DNA double-strand breaks (DSBs) are some of the most toxic DNA lesions, since their inaccurate repair may result in mutations that contribute to cancer onset and progression, and to the development of neurological and immunological disorders ^1^. The formation of DSBs activates a cellular response known as the DNA damage response (DDR), which senses the lesion, signals its presence, and coordinates its repair ^2,3^. Following DSB detection by the MRE11-NBS1-RAD50 (MRN) complex or the single-strand DNA binding protein replication protein A (RPA), apical kinases, such as ataxia-telangiectasia mutated (ATM) and ATM- and Rad3-Related (ATR), are activated and phosphorylate numerous targets, including the histone variant H2AX (named γH2AX). The spreading of γH2AX along the chromosome favors the recruitment of additional DDR proteins, including p53-binding protein (53BP1) and breast cancer 1 (BRCA1), which accumulate in cytologically-detectable DDR foci ^4^. In mammalian cells, DSBs are mainly repaired by re-ligation of the broken DNA ends in a process known as non-homologous end-joining (NHEJ) ^5^. However, during the S/G2 cell-cycle phase, DSBs undergo resection, which directs repair toward homology-based mechanisms ^6^. DNA-end resection is a process initiated by the coordinated action of the MRE11 nuclease within the MRN complex, together with C-terminal binding protein (CtBP) interacting protein (CtIP), and continued by exonuclease 1 (EXO1) ^7^. Resected DNA ends are coated by RPA, which contributes to DDR signaling and undergoes a DNA damage-dependent hyper-phosphorylation ^8^. When complementary sequences are exposed upon resection of both the DSB ends, RAD52 mediates their annealing via a process called single-strand annealing (SSA) resulting in the loss of genetic information ^6^. Alternatively, a homologous sequence located on the sister chromatid or on the homologous chromosome can be used as a template for repair in a process known as homologous recombination (HR) ^9^. The invasion of the homologous sequence is mediated by the recombinase RAD51, whose loading on the ssDNA ends is promoted by breast cancer 2 (BRCA2), which binds BRCA1 through the partner and localizer of BRCA2 (PALB2) ^10,11^. BRCA1, together with its constitutive heterodimer BARD1, is a multifaceted protein with several roles in DDR signaling and repair ^12^. *BRCA1* and *BRCA2* genes are the most frequently mutated genes in breast and ovarian cancer ^13^ and recently-developed drugs, such as poly(ADP-ribose) polymerases (PARP) inhibitors, selectively target cancer cells harboring mutations in these genes ^14^. Among its several functions, BRCA1 promotes DNA-end resection, mainly by counteracting the inhibitory effect of 53BP1 ^15^. Indeed, the HR defect in BRCA1-deficient cells is rescued by the depletion of 53BP1 ^16^.

Recently, a novel role for RNA in the DNA damage signaling and repair has emerged ^17-25^. In particular, we have reported that RNA polymerase II (RNA pol II) is recruited to DSBs, where it synthesizes damage-induced long non-coding RNAs (dilncRNAs) ^17,18^. DilncRNAs are processed to generate DNA damage response RNAs (DDRNAs), which promote DDR signaling ^17,18,21,25^. Similar RNA molecules, named diRNAs, contribute to DSB repair by HR ^22-24^.

It has recently been demonstrated in *Schizosaccharomyces pombe* that DNA:RNA hybrids form at DSBs in a tightly regulated fashion ^26^. Recent reports also suggest that DNA:RNA hybrids may form at DSBs in mammalian cells ^27,28^. However, DNA:RNA hybrid formation at DSBs in mammalian cells has not been investigated in depth yet, nor has any characterization of the molecular mechanisms leading to their formation or metabolism at DSBs been reported. Control of DNA:RNA hybrid levels can be achieved either by avoiding their formation during transcription, or by unwinding or degradation of already-formed hybrids by helicases and RNase H enzymes, respectively. In eukaryotic cells, DNA:RNA hybrids are degraded by RNase H1 and RNase H2, the latter accounting for the majority of RNase H activity in mammalian nuclei. RNase H2 is a heterotrimeric complex composed of a conserved catalytic subunit, called RNase H2A, and auxiliary subunits RNase H2B and RNase H2C, which mediate the interaction with additional proteins ^29^. RNase H2, in addition to its role in removing misincorporated ribonucleotides from genomic DNA ^30,31^, is also responsible for the resolution of DNA:RNA hybrids generated by RNA pol II during transcription ^32,33^. Very recently, RNase H2 was also shown to regulate DNA:RNA hybrid levels at telomeres ^34^. Furthermore, a physiologic role for human RNase H2 was uncovered through the discovery that mutations in any of its subunits cause the Aicardi-Goutieres syndrome (AGS), a neuroinflammatory disease associated with the chronic activation of the immune system in response to an excessive accumulation of aberrant forms of nucleic acids ^35^.

Strong links between HR proteins, RNA, and DNA:RNA hybrids have already been demonstrated and continue to emerge: BRCA1 interacts with several transcription and RNA-processing factors including RNA pol II ^36,37^, recognizes and promotes the processing of miRNA precursors ^38^, and mediates the recruitment of the DNA/RNA helicase SENATAXIN to gene terminators to avoid genome instability induced by DNA:RNA hybrid accumulation ^39^. Depletion of BRCA1 or BRCA2 results in the accumulation of DNA:RNA hybrids globally ^40,41^ and specifically at promoter proximal sites of actively transcribed genes ^42,43^.

In addition, proteins involved in the fanconi anemia (FA) repair pathway are recruited to DNA damage sites via DNA:RNA hybrids to suppress hybrid-associated genomic instability ^44,45^. Altogether these results suggest a yet undefined but emerging intimate relationship between DNA:RNA hybrids and the HR repair pathway.

Herein, we explore the links between RNA, DNA:RNA hybrid formation and metabolism, and HR. We show that dilncRNAs generated at DSBs contribute to the recruitment of the HR proteins BRCA1/BRCA2/RAD51 to DSBs. Specifically, in the S/G2 cell-cycle phase dilncRNAs pair to the resected DNA ends to form DNA:RNA hybrids which are directly recognized by BRCA1 and in this way they facilitate its recruitment to DSBs. Moreover, we demonstrate that RNase H2 is recruited to DSBs in the S/G2 cell-cycle phase and is in a complex with the HR proteins BRCA1/BRCA2/PALB2/RAD51, and that BRCA2 impacts on DNA:RNA hybrid levels by mediating RNase H2 recruitment to DSBs. Combined, our findings suggest a role for DNA:RNA hybrid formation and resolution in the HR process and provide a mechanistic model for the emerging interplay between DNA:RNA hybrids and HR proteins.

## Results

### DNA:RNA hybrids form at resected DNA ends in S/G2-phase cells

Recently, we reported that RNA pol II is recruited to DSBs where it transcribes dilncRNAs bidirectionally starting from exposed DNA ends ^17^. In the same experimental setup where dilncRNAs were characterized, we investigated whether they could form DNA:RNA hybrids. We induced a site-specific DSB by transfecting HeLa cells with the I-PpoI nuclease, whose nuclear localization is induced by 4-hydroxytamoxifen (4-OHT). Upon I-PpoI-mediated DSB generation within the weakly transcribed *DAB1* gene (Supplementary Fig. 1a), we monitored the formation of DNA:RNA hybrids by DNA:RNA hybrids immunoprecipitation (DRIP): briefly, non-crosslinked DNA:RNA hybrids were immunopurified with the specific S9.6 monoclonal antibody and analysed by qPCR. We observed that DSB generation induces the formation of DNA:RNA hybrids peaking at ~1.5kb and up to 3 Kb from both sides of the DSB (Fig. 1a,b), consistently with the already reported dilncRNAs generation upon cut ^17^. Importantly, when cut samples were treated with RNase H, levels of DNA:RNA hybrids strongly decreased, demonstrating the specificity of the DRIP signal (Fig. 1b).

**Fig. 1:**
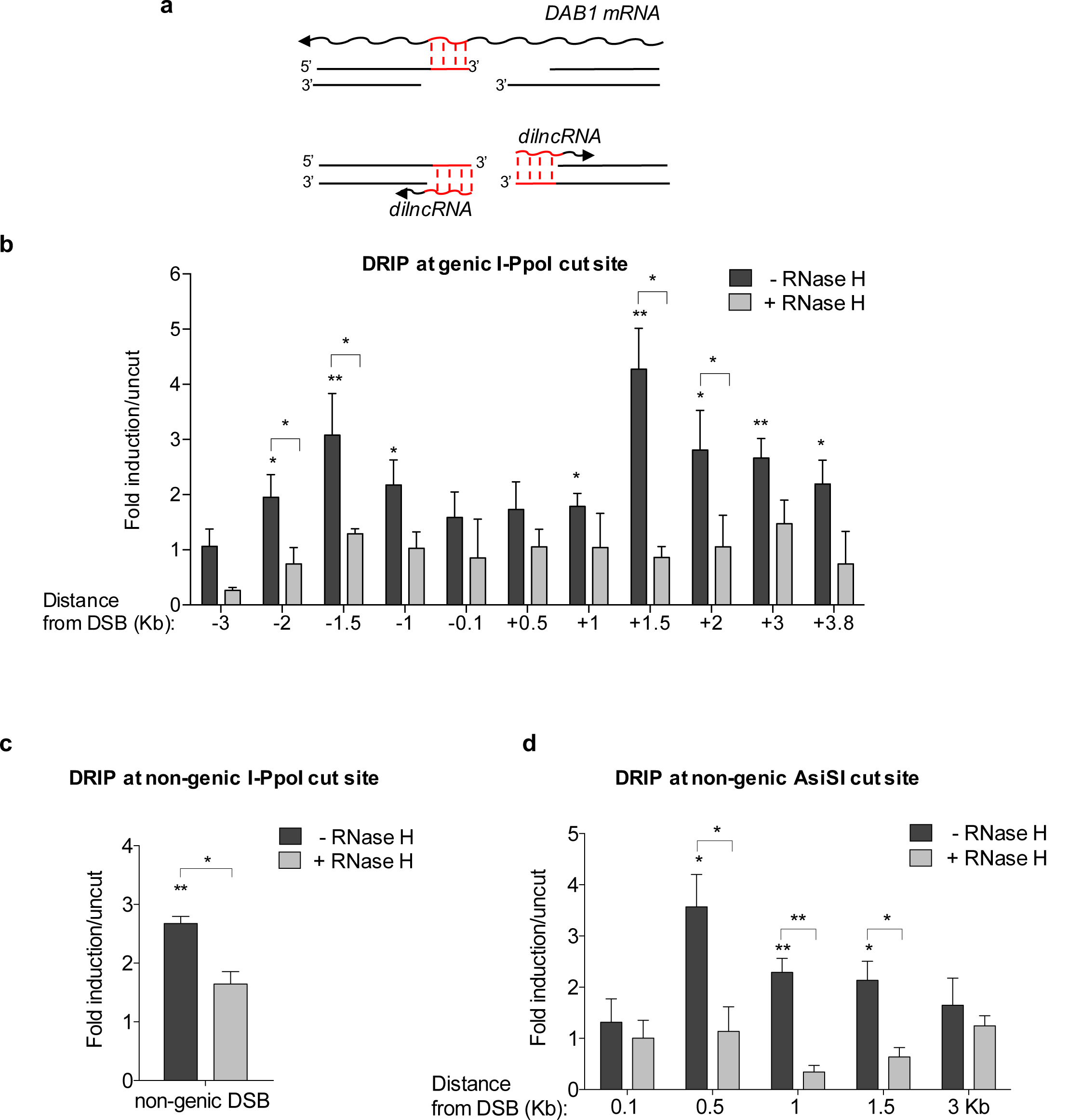
DNA:RNA hybrids form at DBSs. (**a**) Schematic representation of DNA:RNA hybrids (in red) that can be generated upon the hybridization of mRNA (top) or dilncRNAs (bottom) with resected DNA ends at the I-PpoI cut site within *DAB1* gene. (**b**,**c**) DRIP-qPCR analysis at the I-PpoI cut site within a genic (*DAB1* gene) (**b**) or non-genic locus (**c**) in HeLa cells transfected with the I-PpoI nuclease. (**d**) DRIP-qPCR analysis at a non-genic AsiSI cut site in DivA cells. Bar graphs in **b**,**c**, and **d** show fold induction of DNA:RNA hybrid levels in cut samples relative to uncut. RNase H treatment was performed on cut samples to demonstrate specificity of the signal. Error bars represent s.e.m. (*n* ≥ 3 independent experiments). \**P* < 0.05, \*\**P* <0.005 (two-tailed Student’s *t* test).

The accumulation of DNA:RNA hybrids at both sides of the DSB resembled the RNA Pol II-mediated *de novo* bidirectional transcription of dilncRNAs from the DSB ^17^, suggesting that dilncRNAs, rather than pre-existing RNA, such as a mRNA, were generating the observed hybrids (Fig. 1a). To further confirm this observation, we monitored DNA:RNA hybrid accumulation at a DSB within a non-genic I-PpoI target site in HeLa cells (Supplementary Fig. 1a). Indeed, DSB induction by I-PpoI led to DNA:RNA hybrid accumulation also at this site (Fig. 1c), thus supporting the notion that DSBs induce the synthesis of *de novo* transcripts that form DNA:RNA hybrids at DSBs. In order to extend this observation to another non-genic region generated in a different cellular system in which DSBs are induced by a different nuclease, we used DSB inducible via AsiSI (DIvA) U2OS cells (Supplementary Fig. 1a), where nuclear localization of the AsiSI restriction enzyme to the nucleus is induced by 4-OHT to generate DSBs at distinct locations ^46^ (Supplementary Fig. 1b,c). By DRIP-qPCR analyses, we observed DNA:RNA hybrids accumulation at the DSB in a non-genic AsiSI cleavage site (Fig. 1d). Consistently, strand-specific reverse transcription followed by qPCR confirmed dilncRNAs accumulation upon damage at this non-genic AsiSI cleavage site (Supplementary Fig. 1d).

Since dilncRNAs production is dependent on RNA pol II ^17,18^, we tested whether RNA pol II inhibition with 5,6-dichloro-1-β-D-ribofuranosylbenzimidazole (DRB) affected DNA:RNA hybrid generation. DRIP-qPCR at the I-PpoI cleavage site within the *DAB1* gene in HeLa cells treated with DRB, or with vehicle only, prior to DSB induction, revealed that DNA:RNA hybrid accumulation at the damaged site was dependent on RNA pol II (Supplementary Fig. 1e).

Having demonstrated that DNA:RNA hybrids accumulate at I-PpoI- and AsiSI-induced DSBs regardless of the genomic location and that their formation requires RNA pol II activity, we reasoned that, during the S/G2 cell-cycle phase, DNA-end resection and consequent single-stranded DNA generation could provide a suitable DNA substrate for dilncRNA pairing to their resected template DNA and allow hybrids formation (Fig. 1a). We therefore tested whether DNA:RNA hybrid formation is modulated during the cell-cycle by using HeLa-FUCCI cells, which express the fluorescent ubiquitination-based cell-cycle indicators (FUCCI) ^47^. Following I-PpoI expression, we sorted cells into G1- and S/G2-phase populations and we monitored by DRIP-qPCR DNA:RNA hybrid accumulation. Importantly, DNA:RNA hybrids accumulation was analysed at 1.5 Kb on the right from the I-PpoI-induced DSB within the *DAB1* gene, where the resected DNA end could pair only with the newly-synthesised dilncRNA and not with a potentially pre-existing mRNA (Fig. 1a). We observed that, upon DSB induction, DNA:RNA hybrids accumulate preferentially in the S/G2-phase of the cell-cycle (Fig. 2a).

**Fig. 2:**
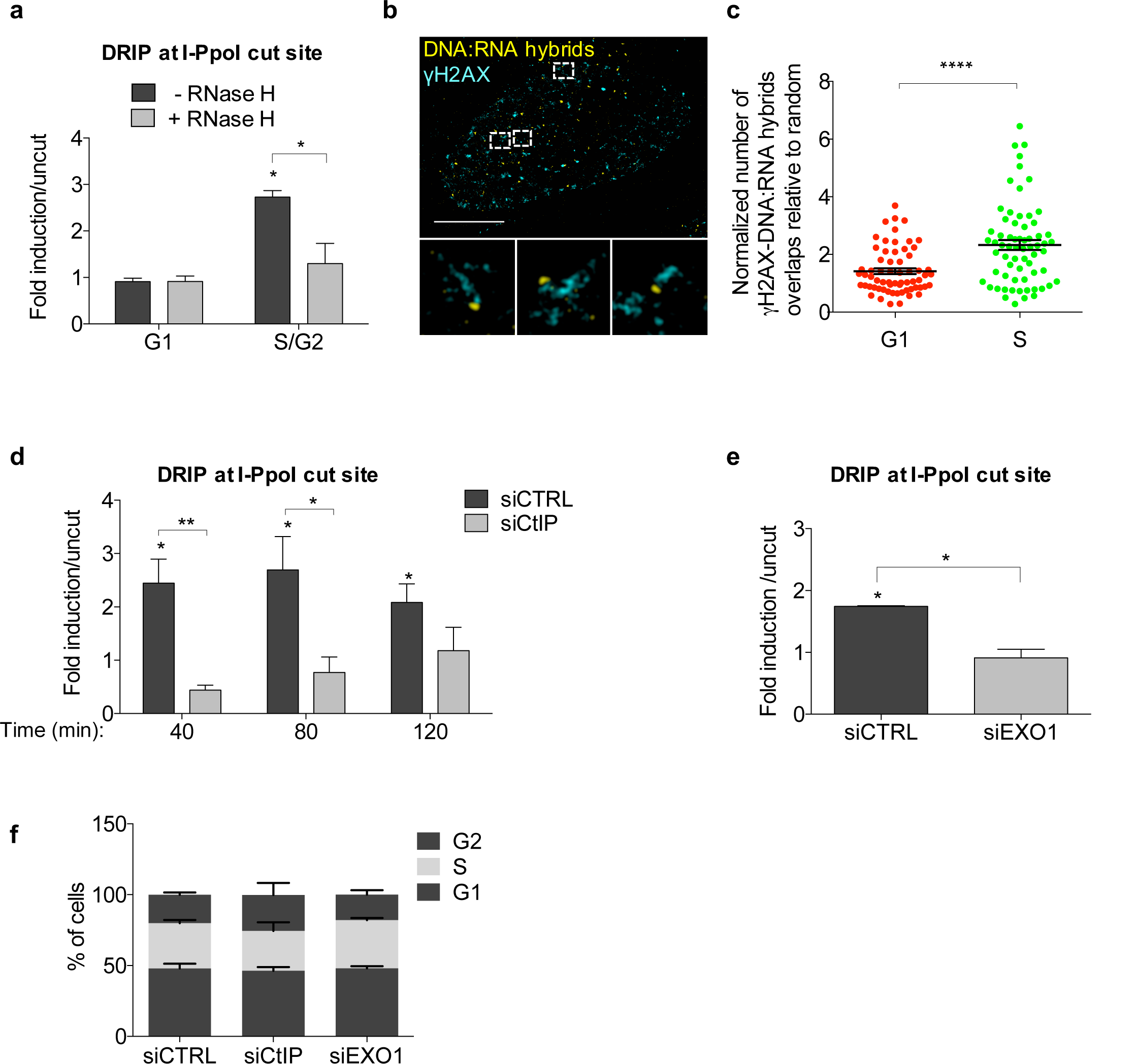
DNA:RNA hybrids form preferentially at resected DNA ends in S/G2-phase cells. (**a**) DRIP-qPCR analysis at 1.5 Kb on the right from the I-PpoI cut site within *DAB1* gene in G1- or S/G2-phase-sorted HeLa-FUCCI cells transfected with the I-PpoI nuclease. The bar graph shows fold induction of DNA:RNA hybrid levels in cut samples relative to uncut. Error bars represent s.e.m. (*n* = 3 independent experiments). (**b**) Representative pictures of super-resolution imaging analysis of γH2AX (cyan) and DNA:RNA hybrids (yellow) co-localization in S-phase synchronized U2OS cells treated with neocarzinostatin (NCS). Scale bar: 5µm. (**c**) Dot plot shows the normalized number of overlaps relative to random of γH2AX and DNA:RNA hybrids signals in G1- or S-phase NCS-treated U2OS cells. Pooled data from *n* = 3 independent experiments are shown. Lines depict the mean±s.e.m. (**d**) DRIP-qPCR analysis at 1.5 Kb on the right from the I-PpoI cut site within *DAB1* gene in cells knocked-down for CtIP at different time points after cut induction. Error bars represent s.e.m. (*n* ≥ 4 independent experiments). (**e**) DRIP-qPCR analysis at 1.5 Kb on the right from the I-PpoI cut site within *DAB1* gene in cells knocked-down for EXO1. Error bars represent s.e.m. (*n* = 2 independent experiments). (**f**) FACS analysis of the cell-cycle profile of cells knocked-down for CtIP or EXO1. Bar graphs represent mean values from *n* = 2 independent experiments. Error bars represent s.e.m. \**P* <0.05, \*\**P* <0.005, \*\*\*\**P* <0.0001 (two-tailed Student’s *t* test).

We next aimed to extend our observations to DSBs formed throughout the genome by an independent approach. To that end, we utilized super-resolution fluorescence microscopy (STORM) and analysed U2OS cells synchronized in G1- or S-phase and treated with the radiomimetic drug neocarzinostatin (NCS). We determined the extent of co-localization between the DDR marker γH2AX and DNA:RNA hybrids detected by S9.6 antibody by quantifying the overlaps of their signals in each cell relative to the calculated number of overlaps present due to random distribution ^48,49^. Importantly, the wider distribution of γH2AX compared to DNA:RNA hybrids was also accounted for in our analysis by this approach. We observed an increased rate of co-localization between γH2AX and DNA:RNA hybrids in S-compared to G1-phase cells (Fig. 2b,c).

Given the preferential DNA:RNA hybrid accumulation at DSBs in the S/G2 cell-cycle phase, we tested the contribution of DNA-end resection to their formation. To this aim, we knocked-down CtIP and we monitored DNA:RNA hybrid accumulation by DRIP-qPCR at 1.5 Kb on the right from the I-PpoI cleavage site within *DAB1* gene, as above, at different time-points after cut induction. We observed that inhibiting resection by knocking-down CtIP prevented DNA:RNA hybrid formation (Fig. 2d and Supplementary Fig. 2a). Impaired DNA:RNA hybrids formation was also observed when extensive DNA end resection was inhibited by EXO1 knock-down (Fig. 2e and Supplementary Fig. 2b). As a control, neither CtIP nor EXO1 knock-down altered cell-cycle phases distribution (Fig. 2f).

Collectively, these results show that DNA:RNA hybrids form at DSBs, as independently demonstrated site-specifically by DRIP analyses at genic and non-genic loci and genome-wide by super-resolution imaging, and that their accumulation requires RNA pol II activity. Moreover, DNA:RNA hybrids formation occurs preferentially during the S/G2 cell-cycle phases and it is favored by DNA-end resection controlled by CtIP and EXO1.

### dilncRNAs contribute to HR proteins recruitment to DSBs and HR-mediated repair

We next studied the impact of transcriptional inhibition on DNA-end resection and on the focal accumulation of HR proteins which are specifically recruited to DSBs in the S/G2 cell-cycle phase. In order to test this, we acutely inhibited RNA pol II activity with α-amanitin or DRB and we simultaneously irradiated (5Gy) HeLa cells – effective RNA pol II inhibition was confirmed by monitoring by RT-qPCR *c-FOS* mRNA level, a specific RNA pol II transcript with a short half-life (Supplementary Fig. 3a,b). DNA-end resection was measured by immunofluorescence analyses of exposed ssDNA through native staining of incorporated BrdU, total RPA and its phosphorylated form (RPA2 pS4/8) – focal signals were quantified in S/G2 cells, as monitored by the S/G2-phase marker cyclin A. Consistently, all three markers revealed proficient DNA-end resection upon RNA pol II inhibition (Fig. 3a,b and Supplementary Fig. 3c). Next, we monitored the impact of transcriptional inhibition by α-amanitin or DRB on the focal accumulation of HR proteins. We observed that both treatments significantly impaired the formation of the foci of BRCA1, BRCA2, and RAD51 (Fig. 3c,d and Supplementary Fig. 3d), this is despite unaltered protein levels (Supplementary Fig. 3e,f).

**Fig. 3:**
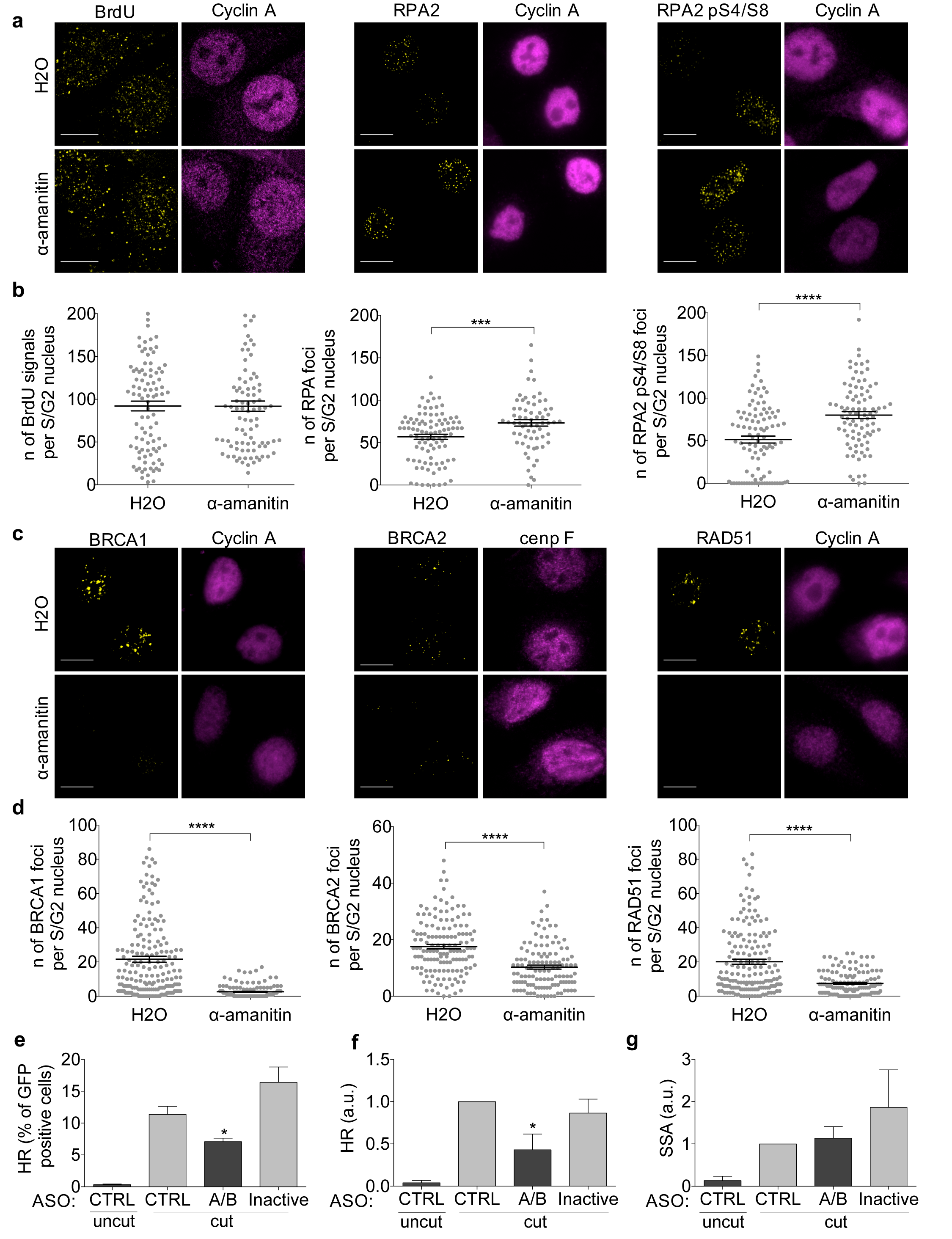
dilncRNAs contribute to HR proteins recruitment to DSBs and HR-mediated repair. (**a**) Representative images of DNA-end resection markers: ssDNA (visualized by BrdU native staining), RPA2, and RPA2 pS4/8 foci, co-stained with cyclin A, as S/G2-phase marker, in irradiated (5Gy) HeLa cells treated with H2O or α-amanitin. Scale bar: 10 µm. (**b**) Dot plots show the number of signals/foci in (**a**). Pooled data from *n* = 3 independent experiments are shown. Lines depict the mean±s.e.m. (**c**) Representative images of BRCA1, BRCA2, and RAD51 foci co-stained with cyclin A or cenp F, as S/G2-phase markers, in irradiated (5Gy) HeLa cells treated with H2O or α-amanitin. Scale bar: 10 µm. (**d**) Dot plots show the number of foci in (**c**). Pooled data from *n* = 3 independent experiments are shown. Lines depict the mean±s.e.m. (**e**,**f**,**g**) DR-GFP cells are treated with control ASO (CTRL), ASOs matching dilncRNAs (A/B), or inactive ASOs. HR efficiency is monitored by FACS analysis of the percentage of GFP positive cells (**e**) or by genomic semi-quantitative PCR (**f**) with primers P1 and P2 that only amplify the recombined GFP sequence generated after HR (see Supplementary Fig. 3g). (**g**) SSA efficiency is assessed by monitoring the 0.8 Kb amplicon generated by genomic semi-quantitative PCR with primers F1 and R2 (see Supplementary Fig. 3g). Bar graphs show the mean of *n* = 3 independent experiments. Error bars represent s.e.m. \**P* <0.05, \*\**P* <0.005, \*\*\**P*<0.001, \*\*\*\**P* <0.0001 (two-tailed Student’s *t* test).

We next sought to test a direct role of dilncRNAs in HR. For this, we employed the DR-GFP reporter cell system ^50^ in which HR between a mutated integrated GFP construct, containing the I-SceI recognition site, and a truncated GFP generates a functional GFP open reading frame (Supplementary Fig. 3g). Following I-SceI induction, HR can be monitored by either the evaluation of GFP expression by FACS analysis in individual cells or, more directly but in bulk, by PCR amplification of the recombined genomic DNA sequence. We impaired dilncRNAs functions by complementary Antisense Oligonucleotides (ASOs) (Supplementary Fig. 3g) – ASOs are modified oligonucleotides widely used to inhibit the function of their target RNAs ^51^ and have been previously used by our group to target dilncRNAs and inhibit DDR ^17,18^. We transfected different sets of ASOs complementary to the predicted dilncRNAs generated at the I-SceI locus and simultaneously induced I-SceI for 72 hours. Both FACS analysis of GFP expression (Fig. 3e) and PCR to detect the recombined genomic locus (Fig. 3f) demonstrated that ASOs matching dilncRNAs reduced HR efficiency, while an ASO matching an unrelated sequence (CTRL) had no impact on HR (Fig. 3e,f). Importantly, the same effective ASOs inactivated by annealing with complementary sequences (Inactive) did not inhibit HR, and all the ASOs left cell-cycle unaltered (Supplementary Fig. 3h). In this same cell system, genomic PCR can also be used to study SSA, a RAD52-dependent but RAD51-independent mechanism that shares with HR the initial DNA-end resection step. We observed that ASOs did not inhibit SSA (Fig. 3g), further indicating that dilncRNAs inactivation impacts the HR process downstream of DNA-end resection.

These results show that dilncRNAs, while not affecting DNA-end resection, contribute to the recruitment of HR proteins to the site of damage and, by doing so, they promote HR.

### DNA:RNA hybrids are directly recognized by BRCA1 *in vitro* and promote its recruitment to DSBs in living cells

Based on our observations that dilncRNAs form DNA:RNA hybrids in S/G2-phase cells and control the recruitment of HR proteins to sites of DNA damage, we sought to test the involvement of DNA:RNA hybrids in the focal accumulation of HR proteins at DSBs. By performing super-resolution imaging and analyzing the extent of co-localization between BRCA1 and DNA:RNA hybrids in NCS-treated U2OS cells synchronized in S-phase, we observed that the few detectable DNA:RNA hybrids often co-localize with BRCA1 in S-phase cells upon damage (Fig. 4a,b).

**Fig. 4:**
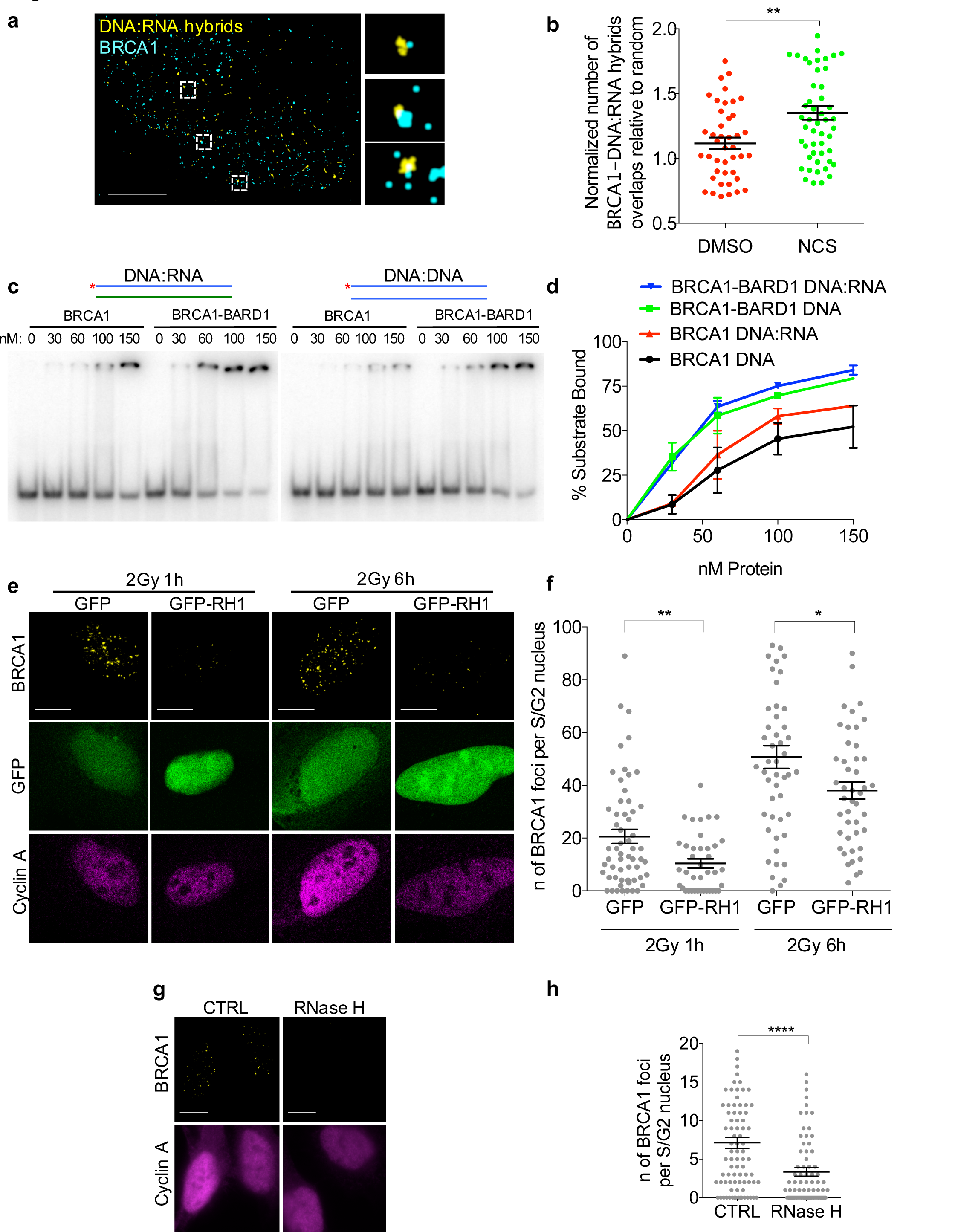
DNA:RNA hybrids are directly recognized by BRCA1 *in vitro* and promote its recruitment to DSBs in living cells. (**a**) Representative pictures of super-resolution imaging analysis of BRCA1 (cyan) and DNA:RNA hybrids (yellow) co-localization in S-phase synchronized NCS-treated U2OS cells. Scale bar: 5µm. (**b**) Dot plot shows the normalized number of overlaps relative to random of BRCA1 and DNA:RNA hybrids signals in S-phase U2OS cells treated with DSMO or NCS. Pooled data from *n* = 3 independent experiments are shown. Lines depict the mean±s.e.m. (**c**) Electrophoretic mobility shift assay (EMSA) of purified recombinant human BRCA1 or BRCA1-BARD1 with end labeled (*) double-stranded DNA or DNA:RNA substrates. (**d**) Graph showing the percentage of protein-bound substrate at respective protein concentrations from *n* = 2 biological replicates. Error bars represent s.e.m. (**e**) Representative images of BRCA1 foci co-stained with cyclin A, as S/G2-phase marker, in irradiated (2Gy) U2OS cells over-expressing GFP or GFP-RNase H1 (GFP-RH1). Scale bar: 10 µm. (**f**) Dot plot shows the number of foci in (**e**). Pooled data from *n* = 3 independent experiments are shown. Lines depict the mean±s.e.m. (**g**) Representative images of BRCA1 foci co-stained with cyclin A, as S/G2-phase marker, in irradiated (2Gy) U2OS cells treated with RNase H prior to fixation. Scale bar: 10 µm. (**h**) Dot plot shows the number of foci in (**g**). Pooled data from *n* = 3 independent experiments are shown. Lines depict the mean±s.e.m. \**P* <0.05, \*\**P* <0.005, \*\*\*\**P* <0.0001 (two-tailed Student’s *t* test).

To test whether BRCA1 can directly recognize DNA:RNA hybrids, we used purified recombinant human BRCA1 or the constitutive BRCA1-BARD1 heterodimer in an electrophoretic mobility shift assay (EMSA) with either DNA duplexes or DNA:RNA hybrids. Radioactively-labelled probes were incubated with the recombinant proteins and separated by electrophoresis on a native polyacrylamide gel. Both BRCA1 alone and BRCA1-BARD1 bound the DNA:RNA hybrid, with an affinity comparable or higher than that for dsDNA (Fig. 4c,d).

Having observed that BRCA1 can bind DNA:RNA hybrids, we tested whether modulation of DNA:RNA hybrids level at DSBs in living cells impacted its recruitment. To this purpose, we monitored BRCA1 foci formation at DSBs in irradiated (2Gy) U2OS cells expressing RNase H1 fused to GFP, or GFP alone as a control. We observed that RNase H1 overexpression impaired ionizing radiation-induced BRCA1 foci formation (Fig. 4e,f), indicating a role for DNA:RNA hybrids in favoring BRCA1 recruitment to DSBs. To rule out any indirect effect of RNase H1 overexpression, we treated irradiated cells with RNase H *in situ*. Briefly, we irradiated U2OS cells (2Gy) and one hour later we gently permeabilized and incubated them with recombinant bacterial RNase H. After 30 minutes, cells were fixed and BRCA1 foci were monitored in S/G2-phase cells. We observed that RNase H treatment reduced the amount of BRCA1 foci (Fig. 4g,h), while not impacting neither on the number of γ-H2AX foci (Supplementary Fig. 4a), nor on DNA-end resection, as determined by RPA foci (Supplementary Fig. 4b).

These results show that DNA:RNA hybrids can be directly recognized by BRCA1 *in vitro*, and in living cells they contribute to BRCA1 recruitment to DSBs.

### RNase H2 is recruited to DSBs during the S/G2 cell-cycle phase

Although DNA:RNA hybrids promote the early steps of HR by favoring BRCA1 loading at DSB, it is known that their excessive accumulation may be detrimental for HR ^28^. This suggests that their levels at DSBs need to be tightly controlled. Since RNase H2 is the major source of RNase H activity in mammalian nuclei ^29^, we tested its recruitment to DSB by performing chromatin immunoprecipitation (ChIP) and assaying for RNase H2A enrichment at the I-SceI cut site in the DR-GFP system. We observed an enrichment of RNase H2A around the I-SceI cleavage site in cut DR-GFP cells relative to uncut cells (Fig. 5a), to an extent similar to that observed for BRCA1 (Fig. 5b), while no enrichment was detected in an unrelated region in cut *versus* uncut (Fig. 5a,b). To further validate this result with a different technique and at multiple genomic sites, we performed immunofluorescence microscopy to detect RNase H2 subunits in irradiated or not irradiated cells. However, such stainings showed a diffuse signal that failed to highlight discrete foci under all the conditions tested (data not shown). To increase the sensitivity and specificity of the signal, we performed proximity ligation assay (PLA) between RNase H2A and γH2AX in not irradiated or irradiated (2Gy) U2OS cells fixed 1 or 6 hours after irradiation. We observed an increase in PLA signals between RNase H2A and γH2AX in irradiated cells (Fig. 5c and Supplementary Fig. 5a), thus suggesting that the two proteins become in close proximity upon irradiation. As a negative control, no signal was detected when only one of the two primary antibodies was used (Supplementary Fig. 5b,c) and, as a reference, a comparable PLA signal was observed between γH2AX and the HR marker RAD51 (Supplementary Fig. 5d). Since damage-induced DNA:RNA hybrids preferentially accumulate in the S/G2 cell-cycle phase, we monitored RNase H2 recruitment to DSBs in irradiated (2Gy) and not irradiated HeLa-FUCCI cells. In this setup, we observed an increase in PLA signals between γH2AX and RNase H2A in S/G2-phase irradiated cells compared to G1 or to S/G2 not irradiated cells (Fig. 5d,e), indicating that RNase H2 localizes to DSBs preferentially in the S/G2 cell-cycle phase. Similar results were also obtained with a different antibody raised against RNase H2A (Supplementary Fig. 5e) or the RNase H2B subunit (Supplementary Fig. 5f). These conclusions were not biased by cell-cycle variations of RNase H2 protein levels or of γH2AX foci, as both the number of γH2AX foci (Supplementary Fig. 5g) and the RNase H2A and RNase H2B pan-nuclear signals (Supplementary Fig. 5h,i) remained unchanged in G1-versus S/G2-phase cells. The observed increased PLA signal in S/G2-compared to G1-phase cells (Fig. 5e), with unaltered levels of both antigens, further demonstrates the specificity of the assay and of the conclusions reached.

**Fig. 5:**
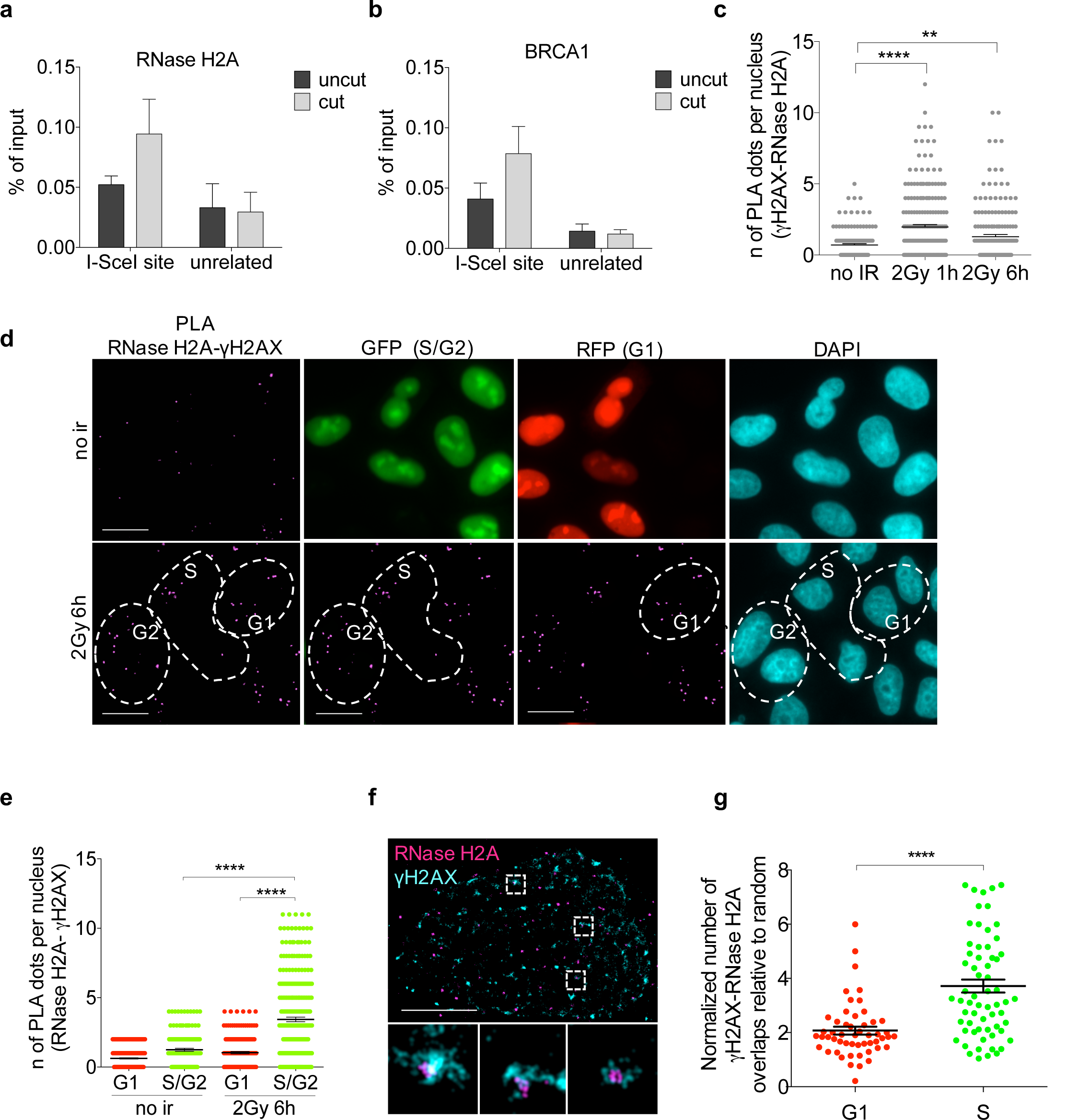
RNase H2 is recruited to DSBs preferentially during the S/G2 phase of the cell cycle. (**a,b**) ChIP of RNase H2A (**a**) or BRCA1 (**b**) at the I-SceI cut site or at an unrelated region in uncut or cut U2OS DR-GFP cells. The bar graph shows the average % of input from *n* = 3 independent experiments. Error bars represent s.e.m. (**c**) Dot plot shows the number of signals per nucleus of PLA between RNase H2A and γH2AX in not irradiated (no ir) or irradiated (2Gy) U2OS cells (shown in Supplementary Fig. 5a). Pooled data from *n* = 3 independent experiments are shown. Lines depict the mean±s.e.m. (**d**) Representative images of PLA between RNase H2A and γH2AX in not irradiated (no ir) or irradiated (2Gy) G1- and S/G2-phase HeLa-FUCCI cells. Scale bar: 10 µm. (**e**) Dot plot shows the number of PLA signals in (**d**). Pooled data from *n* = 3 independent experiments are shown. Lines depict the mean±s.e.m (**f**) Representative pictures of super-resolution imaging analysis of γH2AX (cyan) and RNase H2A (magenta) co-localization in G1- or S-phase synchronized NCS-treated U2OS cells. Scale bar: 5µm. (**g**) Dot plot shows the normalized number of overlaps relative to random of γH2AX and RNase H2A signals in G1- or S-phase cells. Pooled data from *n* = 3 independent experiments are shown. Lines depict the mean±s.e.m. \*\**P* <0.005, \*\*\*\**P* <0.0001 (two-tailed Student’s *t* test).

In order to further extend our observations with an independent approach, we performed super-resolution imaging analysis of γH2AX and RNase H2A co-localization in U2OS cells treated with NCS and we measured the extent of co-localization relative to random events. In agreement with PLA results, we observed that RNase H2 co-localized with γH2AX in NCS-treated S-phase cells (Fig. 5f,g).

Overall, these results consistently indicate that RNase H2 is recruited to DSBs, both induced at a specific locus and genome-wide, preferentially during the S/G2-phase of the cell-cycle.

### BRCA2 is in a complex with RNase H2 and controls DNA:RNA hybrid levels at DSBs

Published reports suggest BRCA2 as a possible regulator of DNA:RNA hybrids cellular levels ^40^. To test whether DNA:RNA hybrid levels at DSBs could be controlled by BRCA2 through RNase H2 recruitment, we performed PLA between γH2AX and RNase H2A in S/G2-phase irradiated (2Gy) HeLa-FUCCI cells knocked-down for BRCA2. We observed that RNase H2A recruitment to DSBs was reduced in cells knocked-down for BRCA2 (Fig. 6a,b and Supplementary Fig. 6a), despite no significant differences in γH2AX foci numbers (Supplementary Fig. 6b). The observed requirement of BRCA2 for RNase H2 recruitment to DSBs prompted us to test whether the two proteins could form a complex. We thus performed immunoprecipitation experiments from cell lysates of irradiated and not irradiated HEK293T cells prepared in the presence of benzonase to degrade all contaminating nucleic acids. We observed that RNase H2A co-immunoprecipitates with BRCA2 and other proteins of the HR machinery, including BRCA1, PALB2, and RAD51, independently of DNA damage induction (Fig. 6c). This interaction was specific within this complex since no interactions with proteins known to be part of other BRCA1 complexes, such as CtIP and RAP80, were observed (Supplementary Fig. 6c).

**Fig. 6:**
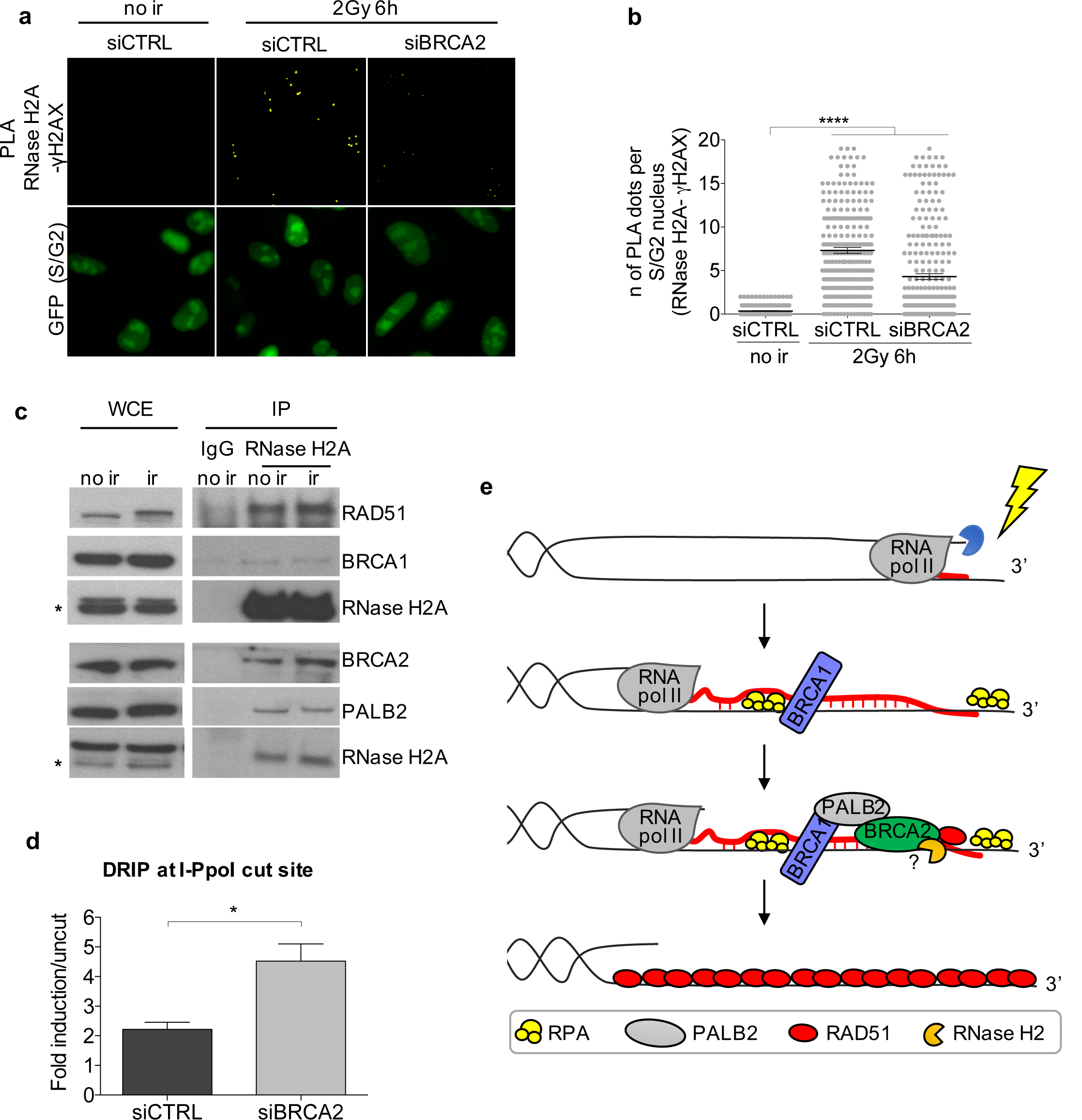
BRCA2 controls DNA:RNA hybrid levels at DSBs via RNase H2 recruitment in S/G2-phase cells. (**a**) Representative images of PLA between RNase H2A and γH2AX in not irradiated (no ir) or irradiated (2Gy) U2OS cells knocked-down for BRCA2. Scale bar: 10 µm. (**b**) Dot plot shows the number of signals per nucleus of PLA between RNase H2A and γH2AX in cells knocked-down for BRCA2. Pooled data from *n* = 3 independent experiments are shown. Lines depict the mean±s.e.m. (**c**) Co-immunoprecipitation of endogenous RNase H2A from not irradiated (no ir) or irradiated 5Gy (ir) HEK293T cell extracts, prepared in the presence of benzonase to avoid contaminant nucleic acids. Asterisks indicate specific band. This experiment was repeated three times independently with similar results. (**d**) DRIP-qPCR at 1.5 Kb on the right from the I-PpoI cut site within *DAB1* gene in S/G2-phase-sorted HeLa-FUCCI cells knocked-down for BRCA2 and transfected with the I-PpoI nuclease. The bar graph shows the average fold induction of cut samples relative to uncut from *n* = 3 independent experiments. Error bars represent s.e.m. (**e**) Model of DNA:RNA hybrid dynamics at DSBs. \**P* <0.05, \*\*\**P*<0.001 (two-tailed Student’s *t* test).

In order to test whether the impaired RNase H2 localization to DSBs upon BRCA2 inactivation resulted in increased DNA:RNA hybrid levels at DSBs, we performed DRIP-qPCR at the I-PpoI site within the *DAB1* gene in S/G2-phase-sorted HeLa-FUCCI cells knocked-down for BRCA2 – sorting of the S/G2-phase cells population was necessary since BRCA2 inactivation affects the cell-cycle (data not shown). DRIP-qPCR analysis revealed a significantly increased accumulation of DNA:RNA hybrids at the DSB in the absence of BRCA2 (Fig. 6d and Supplementary Fig. 6a), indicating that BRCA2, likely via the recruitment of RNase H2, regulates DNA:RNA hybrid levels at DSBs. Interestingly, RAD51 knock-down in the same experimental conditions did not significantly alter DNA:RNA hybrid levels at the tested DSB (Supplementary Fig. 6d,e).

Altogether these results show that RNase H2A is in a specific complex with HR proteins, including BRCA1, PALB2, BRCA2, and RAD51, and that BRCA2 controls DNA:RNA hybrid levels at DSBs by mediating RNase H2 recruitment to DSBs.

## Discussion

We have recently demonstrated that in mammalian cells RNA pol II is recruited to exposed DNA ends upon breakage, where it bidirectionally transcribes RNA species named dilncRNAs ^17^. In the present study, we show that DNA:RNA hybrids form at DSBs most likely upon hybridization of dilncRNAs generated at DSBs with the resected ends of their template DNA in the S/G2-phase of the cell-cycle. In this way, dilncRNAs contribute to the recruitment of key HR proteins to DSBs and, therefore, modulate HR. Specifically, dilncRNAs contribute to the initial BRCA1 recruitment to DSBs by forming DNA:RNA hybrids, which can be directly recognized by BRCA1. However, since an excess of DNA:RNA hybrids could be detrimental for the HR process, we also observe that RNase H2 is in a complex with the HR proteins BRCA1, BRCA2, PALB2, and RAD51 and it is recruited to DSBs in the S/G2 cell-cycle phase most likely via BRCA2 to modulate DNA:RNA hybrids levels. Therefore, our results suggest that while the formation of DNA:RNA hybrids initially promotes BRCA1 recruitment to DSBs, this needs to be tightly controlled by RNase H2 not to prevent the later steps of HR.

Our observation that DNA:RNA hybrids form at DSBs in mammalian cells is in line with recent data in *Schizosaccharomyces pombe* showing DNA:RNA hybrid accumulation at DSBs ^26^ and it is consistent with preliminary evidence in mammalian cells suggesting DNA:RNA hybrid formation at DSBs ^27^. Consistently, the human RNA-unwinding protein DEAD box 1 (DDX1) ^28^ and the DNA/RNA helicase SENATAXIN ^52^ have been shown to localize to DSBs in a transcription- and DNA:RNA hybrids-dependent manner. In particular, the DNA:RNA hybrids-dependent recruitment of DDX1 to DSBs is dependent on DNA-end resection ^53^. In line with this and with data generated in *S. pombe* ^26^, we show that DNA:RNA hybrids form in the S/G2 cell-cycle phase upon exposure of resected DNA ends to the complementary RNAs. Importantly, here we show for the first time that DNA:RNA hybrids form at DSBs in both genic and non-genic regions, thus indicating that the RNA component of these hybrids is unlikely to be be a pre-existing transcript, but rather could be a newly transcribed dilncRNA. Further supporting this observation, we also demonstrate that the presence of resected DNA ends is required for DNA:RNA hybrids accumulation at DSBs. This indicates that DNA:RNA hybrids formation, even in genic regions, cannot only be the result of pairing of the pre-existing mRNA to the template DNA, since it would occur only on one side of the DSB: the one with exposed ssDNA matching the pre-existing transcript (see Fig. 1a). Differently, the observed DNA:RNA hybrids accumulation at both sides of DSBs is only compatible with newly bidirectionally transcribed dilncRNAs pairing with their template resected DNA ends. Interestingly, the reported need for pre-existing transcription to promote RNA-mediated repair in *Saccharomyces cerevisiae* ^54^ is consistent with the reported lack of recruitment of RNA polymerase II to DSB and lack of transcriptional induction in this species ^55^.

Our work also shows that RNA pol II transcription contributes to the focal accumulation of the HR proteins BRCA1, BRCA2, and RAD51 at DSBs, while it does not seem necessary for DNA-end resection. Accordingly, site-specific inactivation of dilncRNAs by complementary ASOs inhibits repair by HR, but does not affect SSA, which requires extensive DNA ends resection but differs from HR in the subsequent steps. The observation that transcriptional inhibition does not reduce DNA-end resection while impairing BRCA1 foci formation was unexpected. However, it can be explained by considering the concomitant reduced 53BP1 foci formation upon transcriptional inhibition or ASO treatment ^17^. Indeed, since BRCA1 is required to oppose the inhibitory effect of 53BP1 on DNA-end resection, in the absence of 53BP1, as upon RNA pol II inhibition or ASO treatment, BRCA1 may become dispensable for this process ^15^. This could also explain the observed stronger impact of transcriptional inhibition on BRCA1 recruitment to DSBs compared to BRCA2 and RAD51. Notably, the moderate increase of DNA-end resection observed upon transcriptional inhibition may be caused either by a higher efficiency of the resection process, or, more intriguingly and consistently with our model, by an increased availability of single-stranded DNA for RPA binding in the absence of a competing complementary RNA paired to the resected DNA end.

At DSBs BRCA1 can be detected co-localizing with DNA:RNA hybrids and these hybrids contribute to its recruitment to DSBs, as shown by the reduced number of ionizing-radiation induced BRCA1 foci upon RNase H1 overexpression or treatment *in situ* with RNase H. In particular, we also provide the first direct evidence that both the purified recombinant human BRCA1 and the constitutive BRCA1-BARD1 heterodimer can bind DNA:RNA hybrids *in vitro*, with an affinity similar to the well-characterized dsDNA substrate. This result provides evidence in support of a direct interaction between BRCA1 and DNA:RNA hybrids, and it is consistent with the observed DNA:RNA hybrid-dependent BRCA1 recruitment to gene termination sites in living cells ^39^.

Recent evidence shows that excessive amounts of DNA:RNA hybrids at DSBs may dampen repair by HR, as demonstrated by impaired HR efficiency in the absence of the RNA-unwinding protein DDX1 ^28^. In line with these results, in human and *Drosophila* cells, the catalytic component of the RNA exosome, which contributes to RNA degradation, localizes to DSBs and its activity is required for RAD51 loading ^56^. In *S. pombe*, only controlled levels of DNA:RNA hybrids at DSBs facilitate repair ^26^. Similarly, a precise level of DNA:RNA hybrids is required to guarantee the proper length of mammalian telomeres that elongate through an HR-based pathway named Alternative Lengthening of Telomeres (ALT) ^57^. In mammalian cells, emerging links between HR proteins and DNA:RNA hybrid levels have recently been described and support our conclusions. DNA:RNA hybrids accumulate globally in cells lacking BRCA1 or BRCA2 ^40,41^. Additionally, proteins that, together with BRCA2, control the FA repair pathway localize to DNA damage sites via DNA:RNA hybrids ^44,45^. However, until now, no mechanisms explaining this emerging link between proteins controlling HR or DNA:RNA hybrid metabolism have been proposed. Here, we provide the first evidence that RNase H2, the main protein responsible for DNA:RNA hybrid degradation in mammalian nuclei ^29^, interacts with the HR machinery components, including BRCA1, BRCA2, PALB2, and RAD51, and that BRCA2 contributes to its recruitment to DSBs specifically during the S/G2 cell-cycle phase. Indeed, BRCA2 inactivation boosts DNA:RNA hybrid levels at DSBs. This observation is not only consistent, but it could actually mechanistically explain the increased DNA:RNA hybrid levels observed in BRCA2-depleted cells by others ^40^. In light of our conclusions, it will be interesting to study the DNA:RNA hybrid levels at DSBs in conditions in which RNase H2 is mutated, as in AGS patients.

In summary, we propose a model (Fig. 6e) in which DNA:RNA hybrids form upon hybridization of dilncRNAs with resected DNA ends generated during the S/G2 cell-cycle phase. DNA:RNA hybrids are initially recognized by BRCA1 and subsequently, BRCA2-mediated recruitment of RNase H2 induces their degradation, thus ensuring efficient HR-mediated repair.

## Author contributions

M.A. performed immunoprecipitation experiments; D.W. performed super-resolution imaging analyses; S.H. performed the EMSA with purified recombinant proteins. G.D and C.J-W. performed DR-GFP experiments. G.D. generated all remaining data and wrote the manuscript; F.d’A.d.F. supervised the project and revised the manuscript; all authors edited the manuscript.

## Acknowledgements

We thank G. Legube (Centre de Biologie Intégrative, Toulouse, France), M. Kastan (Duke Cancer Institute, Durham, USA), N. Proudfoot (Sir william dunn school of pathology University of Oxford), and B. Xia (The Cancer Institute of New Jersey, University of Medicine and Dentistry of New Jersey, USA) for reagents and all F.d’A.d.F. group members for reading the manuscript, support and constant discussions. G.D. was supported by Fondazione Italiana Ricerca Sul Cancro (Application No. 15050). C.J-W is supported by Fondazione Italiana Ricerca Sul Cancro (Application No. 19589). Work in the Rothenberg laboratory is supported by grants from the NIH (CA187612, GM108119) and the American Cancer Society (RSG DMC-16-241-01-DMC). Work in Petr Cejka’s laboratory is supported by European Research Council (681630) and the Swiss National Science Foundation (31003A_175444). Work in Fabrizio d’Adda di Fagagna’s laboratory is supported by the Associazione Italiana per la Ricerca sul Cancro, AIRC (application 12971), Cariplo Foundation (grant 2010.0818 and 2014-0812), Fondazione Telethon (GGP12059), Association for International Cancer Research (AICR-Worldwide Cancer Research Rif. N. 14-1331), Progetti di Ricerca di Interesse Nazionale (PRIN) 2010–2011, the Italian Ministry of Education Universities and Research EPIGEN Project, a European Research Council advanced grant (322726) and AriSLA (project ‘DDRNA and ALS’).

## Competing interests

The authors declare no competing financial interests.

## Methods

### Cell culture

All the cell lines were grown under standard tissue culture conditions (37°C, 5% CO_2_). HeLa were grown in Minimum Essential Medium (MEM) (Biowest/Gibco) supplemented with 10% fetal bovine serum (FBS), 1% L-glutamine, non-essential amino acids (10mM for each aa) and 1mM NaPyruvate; U2OS cells were grown in McCoy (Gibco) supplemented with 10% FBS, 1% L-glutamine, and 1% penicillin/streptomycin. HeLa-FUCCI (RIKEN BioResource Center cell bank) ^47^ and doxycycline-inducible I-SceI/DR-GFP (TRI-DR-U2OS) (kind gift from P. Oberdoerffer) were cultured in Dulbecco’s Modified Eagle’s medium (DMEM) (Lonza) supplemented with 10% FBS, 1% L-glutamine, and 1% penicillin/streptomycin. For TRI-DR-U2OS cells, I-SceI expression was induced by adding 5 µg/ml doxycycline to the cell medium. DIvA cells (AsiSI-ER-U20S) (kind gift from G. Legube) were cultured in DMEM without phenol red (Gibco) supplemented with 10% FBS, 1% L-Glutamine, 1% pyruvate, 2.5% HEPES, 1% penicillin/streptomycin, and 1 µg/ml puromycin. AsiSI-dependent DSBs induction was obtained by treating the cells with 300 nM 4 hydroxytamoxifen (4OHT) (Sigma-Aldrich) for 4 h.

U2OS cell synchronization for super-resolution imaging experiments was obtained by serum starvation. Briefly, cells were plated on glass coverslips for 24 h. G0/G1 phase synchronization was achieved by replacing complete medium with serum free medium for 72 h. A mid-S phase cell population was obtained after 16 h release into complete medium. Double strand breaks (DSBs) were generated by using the radiomimetic drug Neocarzinostatin (NCS) (Sigma-Aldrich).

All cell lines are tested for mycoplasma by PCR and by a biochemical test (MycoAlert, Lonza).

Ionizing radiation was induced by a high-voltage X-ray generator tube (Faxitron X-Ray Corporation).

### Plasmids, siRNA, and ASOs transfection

2 µg of GFP-RNase H1 plasmid (kind gift from N. Proudfoot) and its related control was transfected with Lipofectamine 2000 (Life Technologies) and experiments were performed 24 h after transfection.

1 µg of mammalian ER-I-PpoI (kind gift from M. Kastan) was transfected with Lipofectamine 2000. 24 h after transfection, nuclear translocation of ER-I-PpoI was induced by adding 4-OHT (Sigma-Aldrich) at 2 µM final concentration for the indicated time. When I-PpoI was transfected, medium without phenol red was used to avoid leakiness in the expression of the plasmids.

RNA interference was achieved by transfecting 5-20 nM siRNAs with Lipofectamine RNAiMAX transfection reagent (Life Technologies) for 48 hours. For DNA:RNA hybrids detection upon EXO1, BRCA2, or RAD51 knock-down, cells were seeded and transfected with siRNAs in parallel. I-PpoI was transfected the day after. For DNA:RNA hybrids detection upon CtIP knock-down, cells were transfected with siCtIP for 48 h, then replated and transfected with I-PpoI. 24 h later I-PpoI expression was induced by 4-OHT. Sequences of the siRNA used are listed in table 1.

Antisense oligonucleotides (ASOs), mixmers containing locked nucleic acid oligonucleotides with a fully phosphorothioate backbone (Exiqon), were transfected with Lipofectamine RNAiMAX transfection reagent (Life Technologies) at a final concentration of 20 nM. Before transfection, ASO solution was incubated at 95°C for 5 min and chilled on ice for 5 min, to prevent the formation of secondary structures of the oligonucleotides.

For DR-GFP experiments, concomitantly with ASOs transfection, I-SceI expression was induced by adding doxycycline to the cell media and HR was evaluated 72 h after, as described in the section “DR-GFP reporter assay”. ASO sequences are listed in table 2.

### Inhibition of RNA polymerase II transcription

RNA polymerase II transcription was inhibited by treatment with α-amanitin (50 µg/ml) or 5,6-dichloro-1-β-D-ribofuranosylbenzimidazole (DRB, 50 µM), respectively dissolved in deionized water and Dimethyl sulfoxide (DMSO). Immediately after adding the drugs, HeLa cells were irradiated (5Gy) and then fixed 6 h later. For α-amanitin treatment, before adding the drug to the medium, cells were mildly permeabilized with 2% Tween 20 in PBS 1X for 10 min at RT. RT-qPCR analysis of the levels of c-FOS, a short-lived RNA specifically transcribed by RNA pol II, was used to monitor the efficacy and specificity of the drugs.

### RNase H treatment

U2OS cells were plated on coverslips and irradiated (2Gy) and RNase H treatment was performed according to a published protocol ^53^. Next, coverslips were washed twice in PBS 1X and fixed and stained as described in the “immunofluorescence” section.

### RNA extraction

Total RNA from cultured cells was extracted with Maxwell^®^ RSC simplyRNA Tissue Kit with the Maxwell^®^ RSC Instrument (Promega), according to manufacturer instructions. For dilncRNAs detection, chromatin-bound RNA was extracted as follows. Cells were fractionated according to a published protocol ^58^. The obtained chromatin fraction was treated with 50 U of Turbo DNase (Ambion) for 10 min at 37 °C and then digested with with 200 µg of Proteinase K (Roche) for 10 min at 37 °C. The RNA was then purified with Maxwell^®^ RSC simplyRNA Tissue Kit.

### Standard RT–qPCR and strand-specific RT–qPCR

For standard RT–qPCR, cDNA was obtained using the SuperScript VILO Reverse Transcriptase (Life Technologies), according to manufacturer instructions. Roche SYBR Green-based RT–qPCR experiments were performed on a Roche LightCycler 480 machine. RPPO was used as normalizer.

For dilncRNAs detection, chromatin-bound RNA was retro-transcribed using the Superscript IV First Strand cDNA synthesis kit (Invitrogen) with strand-specific primers. Expression of dilncRNAs was determined by RT–qPCR using EvaGreen Supermix (Bio-Rad).

### Immunofluorescence and imaging analysis

Cells were fixed in 4% paraformaldehyde (PFA) for 10 min at RT. For BRCA2 staining cells were fixed in ice-cold methanol for 10 min. Immunofluorescence was performed as described previously ^54^. Secondary antibodies used were: goat anti-rabbit or anti-mouse Alexa 488 IgG (Life Technologies); donkey anti-mouse or anti-rabbit Cy3 IgG (Jackson Immuno Research), donkey anti-mouse or anti-rabbit Alexa 647 IgG (Life Technologies). For BrdU native staining cells were incubated with BrdU (Sigma-Aldrich, 10 µg/ml) for 24 h.

Immunofluorescence images were acquired using a widefield Olympus Biosystems Microscope BX71 and the MetaMorph software (Soft Imaging System GmbH). Confocal sections were obtained with a Leica TCS SP2 or AOBS confocal laser microscope by sequential scanning. Comparative immunofluorescence analyses were performed in parallel with identical acquisition parameters. Images were analysed by CellProfiler 2.1.1 software ^59^.

### Super-Resolution (SR) imaging

SR experiments were performed on U2OS cells seeded on coverslips and synchronized as described in “Cell culture” section. Upon DNA damage induction, cells were pre-extracted at RT for 3 min in CSK buffer (10 mM Hepes, 300 mM sucrose, 100 mM NaCl, 3 mM MgCl_2_, and 0.5% Triton X-100, pH 7.4) and fixed for 15 min in PFA (3.7% from 32% EM grade, Electron Microscopy Sciences, 15714) and glutaraldehyde (0.3% from 70% EM grade, Sigma-Aldrich, G7776) in PBS. Blocking was performed in blocking buffer (2% glycine, 2% BSA, 0.2% gelatin, and 50 mM NH_4_Cl in PBS) for 1 hour at RT. Primary and secondary antibodies used are listed in table 4. Immediately before imaging analysis, coverslips were mounted onto a microscope microfluidics chamber and freshly prepared SR imaging buffer, comprising an oxygen scavenging system including 1 mg/ml glucose oxidase (SigmaAldrich, G2133), 0.02 mg/ml catalase (SigmaAldrich, C3155), 10% glucose (SigmaAldrich, G8270) and 100 mM mercaptoethylamine (Fisher Scientific, BP2664100) in PBS, was added to the imaging chamber. Images were acquired with a custom-built SR microscope based on a Leica DMI 3000 inverted microscope. For each field 2000 sequential frames of single molecule emissions at 40 Hz were collected and imaged on an electron-multiplying charged coupled device (EMCCD, Andor) using Solis software (Andor). Each raw image stack was processed for single molecule localization and rendered using 20 nm pixels via rapidSTORM or QuickPALM. Monte Carlo simulations were used to randomly rearrange the clusters within an ROI to calculate a baseline level of random co-localization. Using this approach, 20 random simulations were generated for each nucleus in a pair-wise fashion, examining Red/Green, Red/Blue, and Green/Blue overlap. The total number of overlaps detected in each nucleus (typically 15-100) was normalized to the determined random level of overlap by dividing the number of real overlaps by the average number of overlaps in the same randomly-simulated nucleus. For display purposes, images were smoothed by applying a Gaussian blur filter and colors were thresholded for optimal production of a clear picture of the single foci.

### Proximity Ligation Assay (PLA)

Cells were labeled according to the manufacturer’s instructions (Sigma). Briefly, cells were fixed as described in the section “immunofluoresce and imaging analysis” and incubated with primary antibodies overnight at 4°C. PLA probes (secondary antibodies conjugated with oligonucleotides) were added to the samples. After ligation of the oligonucleotide probes in close proximity (<40 nm), fluorescently labeled oligonucleotides were added together with a DNA polymerase to generate a signal detectable by a fluorescence microscope. Images were acquired using a widefield Olympus Biosystems Microscope BX71 and the MetaMorph software (Soft Imaging System GmbH) and quantification of PLA dots was performed with the automated image-analysis software CellProfiler 2.1.1.

### DR-GFP reporter assay

I-SceI expression was induced by adding 5 µg/ml doxycycline and 72 h later the HR efficiency was determined by quantifying GFP-positive cells (product of successful HR) by flow cytometry, as described in the section “Fluorescence-activated cell sorting (FACS)”. A PCR method was also used to monitor HR and SSA. DNA was extracted with the DNeasy Blood & Tisue Kit (Qiagen) and PCR was performed with the GoTaq^®^ DNA Polymerase (Promega). HR was monitored by measuring the intensity of the amplicon generated by the primers P1 and P2 ^60,61^. SSA was monitored by measuring the intensity of the amplicon generated by amplification with primer F and R2 ^60,61^. Both the measurements were normalized on the actin amplicon. Sequences of the primers are listed in table 3.

### Immunoblotting

Cells were lysed in Laemmli sample buffer (2% sodium dodecyl sulphate (SDS), 10% glycerol, 60 mM Tris-HCl pH 6.8). Protein concentration was determined by the biochemical Lowry protein assay and the desired amount of protein was mixed with bromophenol blue and dithiothreitol (DTT), heated at 95°C for 5 min, and resolved by SDS polyacrylamide gel electrophoresis (SDS-PAGE). After transferring on a nitrocellulose membrane (0,45mm) (400 mA; 1 h) in transfer buffer (25 mM Tris-HCl, 0.2 M Glycine, 20% methanol), membranes were blocked with 5% milk in TBS-T buffer (Tween20 0.1%) for 1 hour at RT and next incubated overnight at 4^o^C with primary antibodies diluted in in 5% milk in TBS-T (see table 4). Next, the membranes were washed with TBS-T 3 times for 10 min and incubated with secondary horseradish peroxidase (HRP)-conjugated antibodies (Bio-rad) diluted in 5% milk in TBS-T. After 3 more washes with TBS-T, HRP activity was detected using a Chemidoc imaging system (Bio-Rad) machine after adding the substrate for the enhanced chemiluminescent reaction ECL (GE Healthcare).

### Immunoprecipitation

HEK293T cells were collected and lysed in TEB150 lysis buffer (50 mM HEPES pH 7.4, 150 mM NaCl, 2 mM MgCl_2_, 5 mM EGTA pH 8, 1 mM DTT, 0.5% Triton X-100, 10% glycerol, protease inhibitor cocktail set III (Calbiochem) and Benzonase 1:1000 (Sigma) for 45 min at 4^o^C. Usually, 1 mg of the protein lysate was used per each immunoprecipitation in a reaction volume of 500 l. As input, 1% of the immnoprecipitation reaction was collected and denatured in the sample buffer (50 mM Tris-HCl pH 6.8; 2% SDS; 10% glycerol; 12.5 mM EDTA; 0.02 % bromophenol blue; 100 M DTT) for 10 min at 95^o^C. Unspecific binding of proteins to the beads was reduced by incubating samples with Protein G beads (50 μl) (Zymed Laboratories) for 1 hour at 4^o^C (pre-clearing). Binding reactions were performed overnight at 4 °C and were followed by addition of protein G–Sepharose beads for 2 h. After 3-6 washes with lysis buffer, immunoprecipitated proteins were released by the addition of sample buffer and incubation at 95^o^C for 10 min.

### Chromatin Immunoprecipitation (ChIP)

ChIP was performed as described previously ^62^. Briefly cells were cross-linked and chromatin was sonicated with a Focused-Ultrasonicator Covaris to obtained fragments of ~500 bp. 50 μg of chromatin were used per sample. DNA was cleaned up by QIAquick PCR purification column (Qiagen) according to the manufacturer’s instructions and Roche SYBR Green-based qPCR experiments were performed on a Roche LightCycler 480 machine (see table 3 for primers sequence).

### DNA:RNA hybrids immunoprecipitation (DRIP)

DRIP was performed following a published protocol ^40^. Briefly, DNA was extracted gently with phenol:chloroform:isoamyl alcohol and digested with *Hind*III, *Eco*RI, *Bsr*GI, *Xba*I and *Ssp*I. After purification from restriction enzymes, half of the DNA was treated overnight with RNase H (NEB). In the meantime, serum-free medium containing the S9.6 antibody (kind gift from M. Foiani) was mixed with protein A and protein G Dynabeads (Invitrogen) and incubated on a rotating wheel overnight at 4°C. After a further purification step, 4 µg of DNA was used for each IP. After elution from the beads, DNA was cleaned up with QIAquick PCR purification column (Qiagen) according to the manufacturer’s instructions. The indicated regions were amplified by Roche SYBR Green-based qPCR on a Roche LightCycler 480 machine (see table 3 for primers sequence). The signal intensity plotted is the relative abundance of DNA–RNA hybrid immunoprecipitated in each region, normalized to input values.

### Fluorescence-activated cell sorting (FACS)

For GFP analysis, 10^6^ cells were fixed in 1% formaldehyde for 20 min on ice. Next, cells were washed in PBS with 1% BSA and fixed in 75% ethanol. Fixed cells were washed again in PBS with 1% BSA and stained with propidium iodide (PI) (Sigma-Aldrich, 50 µg/ml) in PBS supplemented with RNase A (Sigma-Aldrich, 250 µg/ml).

For cell cycle analysis, 10^6^ cells were directly fixed in 75% ethanol, as described above. Samples were acquired on an Attune NxT machine and analyzed with FlowJo_V10 software. At least 10^4^ events were analyzed per sample.

For sorting of HeLa-FUCCI cells, cells were collected in PBS with 2% FBS and the G1 and S/G2 population were sorted with a MofloAstrios (Beckman Coulter) in PBS supplemented with RNaseOUT (Thermo Fisher). Sorted samples were processed for DRIP as described in the section “DNA:RNA hybrids immunoprecipitation (DRIP)”.

### Cloning, expression and purification of recombinant proteins

Recombinant BRCA1 was expressed and purified as a complex in *Sf*9 cells by co-infection with baculoviruses prepared from individual pFastBac1 plasmids pFB-2xMBP-BRCA1-10xHis. Bacmids, primary and secondary baculoviruses were obtained using standard procedures according to manufacturer’s instructions (Bac-to-Bac, Life Technologies). *Sf*9 cells were transfected using a Trans-IT insect reagent (Mirus Bio).

For the large-scale BRCA1 expression and purification, *Sf*9 cells were seeded at 0.5×106 per ml and infected 16 h later with recombinant baculoviruses expressing pFB-2xMBP-BRCA1-10xHis. The infected cells were incubated in suspension at 27°C for 52 h with constant agitation. All purification steps were carried out at 4°C or on ice. The *Sf*9 cell pellets were resuspended in 3 volumes of lysis buffer (Tris-HCl, pH 7.5, 50 mM; Dithiothreitol (DTT), 1 mM; ethylenediaminetetraacetic acid (EDTA), 1 mM; Protease inhibitory cocktail, Sigma P8340, 1:400; phenylmethylsulfonyl fluoride (PMSF), 1 mM; leupeptin, 30 µg/ml; NP40, 0.5%) for 20 min with continuous stirring. Glycerol was added to 16% (v/v) concentration. Next, 5 M NaCl was added slowly to reach a final concentration of 305 mM. The cell suspension was further incubated for 30 min with continuous stirring, centrifuged at 57′800 g for 30 min to obtain soluble extract. Pre-equilibrated amylose resin (New England Biolabs) was added to the cleared soluble extract and incubated for 1 h with continuous mixing. The resin was then collected by centrifugation at 2′000 g for 2 min and washed extensively batch wise as well as on disposable columns (Thermo Scientific) with wash buffer (Tris-HCl, pH 7.5, 50 mM; β-mercaptoethanol, 2 mM; NaCl, 300 mM; glycerol, 10%; PMSF, 1 mM; NP40, 0.5%). Protein was eluted with wash buffer containing 10 mM maltose (Sigma). The eluates were further treated with PreScission protease for 90 min to cleave off the maltose binding protein affinity tag (MBP). The sample was then supplemented with 20 mM imidazole and further incubated with pre-equilibrated Ni-NTA agarose resin (Qiagen) for 1 h. The Ni-NTA resin was transferred on a disposable column and washed extensively with Ni-NTA wash buffer (Tris-HCl, pH 7.5, 50 mM; β-mercaptoethanol, 2 mM; NaCl, 1 M; glycerol, 10%; PMSF, 1 mM; imidazole, 20 mM). Prior to elution, the protein was washed once with the same NiNTA wash buffer listed above but with only 150 mM NaCl. Pooled fractions were stored at −80°C.

### Electrophoretic mobility shift assay

The 50 bp-long dsDNA substrate was prepared by annealing oligonucleotides X12-3 (5’ GACGTCATAGACGATTACATTGCTAGGACATGCTGTCTAGAGACTATCGC3′) and X12-4C (5’ GCGATAGTCTCTAGACAGCATGTCCTAGCAATGTAATCGTCTATGACGTC 3′) as previously described ^63^. The 50 bp-long DNA:RNA hybrid substrate was prepared by annealing X12-3 DNA and X12-4C RNA. BRCA1 was incubated for 30 min at 37º C with 10 nM dsDNA or DNA:RNA hybrid substrate in a reaction buffer containing Tris-HCl pH 7.5, 25 mM; KCl, 90 mM; EDTA pH 8.0, 1 mM; DTT, 1 mM; BSA (NEB), 0.1 mg/ml; RNaseOUT (Invitrogen), 1X. Loading dye (5 µL; 50% glycerol; Tris-HCl pH 7.5, 20 mM; EDTA, 0.5 mM; bromophenol blue) was added to reactions and products were separated by 6% non-denaturing polyacrylamide gel electrophoresis at 4 °C. The gels were dried on 17CHR filter paper (Whatman), exposed to storage phosphor screens, and scanned by a Typhoon Phosphor imager (FLA 9500, GE Healthcare). Quantification of protein-bound substrates was done using Imagequant software.

### Statistical analysis

Statistical analysis was performed with the unpaired two tailed Student’s t-test and represented as a mean ± standard error of the mean (s.e.m.). Asterisk in the figures indicates p value: \**P* < 0.05, \*\**P* < 0.005, \*\*\**P*<0.001, \*\*\*\**P* < 0.0001.

